# Adaptive excursions across the drift barrier

**DOI:** 10.64898/2026.09.25.754336

**Authors:** Stephen P. De Lisle

**Affiliations:** Department of Environmental and Life Sciences, Karlstad University, Universitetsgatan 2, SE-651 88, Karlstad, Sweden

**Keywords:** drift barrier, phenotypic selection, quantitative genetics

## Abstract

Random genetic drift, rather than selection, is expected to determine the dynamics of alleles with fitness effects that fall below the reciprocal of effective population size (*N_e_*). This is the drift barrier. The drift barrier hypothesis suggests that adaptation may be hopelessly constrained in organisms with low *N_e_*, with much of the complexity of life reflecting a byproduct of ineffective natural selection. Yet decades of empirical research in evolutionary ecology provides substantial evidence of adaptation in finite metazoan populations. Here I explore the extent to which the drift barrier may, or may not, constrain contemporary adaptive divergence under directional selection in the wild. Compensatory directional selection on polygenic traits occurring after the fixation of a deleterious allele will be twice as efficient compared to the initial action of stabilizing selection. Consistent with this result, meta analysis reveals a negative relationship between contemporary estimates of *N_e_* and the strength of directional selection in natural populations. Analysis of empirical phenotypic selection gradients from wild metazoans further reveals that directional selection is often strong enough to overcome the constraints imposed by finite population size, across a range of genetic architectures. Moreover, on a multi-peak adaptive landscape, the probability of a peak shift between adaptive zones increases exponentially as *N_e_* declines, with observed values of contemporary *N_e_* typically being well within the range needed for peak shift models to be a viable explanation of diversity. Together, these results suggest that widespread adaptation in metazoans is not inconsistent with the fundamental constraints on perfection imposed by drift.

## Introduction

Natural selection is limited by both genetics and demography. For natural selection to result in evolutionary change, there must be a genetic basis to phenotypic variation (Fisher, 1930; Lande and Arnold, 1983). Furthermore, selection on genetic variants must be stronger than the stochastic force of genetic drift, which is a function of population size, if selection is to result in change in the allele frequency at a given locus (Kimura, 1983; Ohta, 1973). While these two related phenomenon are widely recognized as constraints on the efficacy of natural selection, the first, that of constraints imposed by a lack of genetic variation, is typically thought to not immediately curtail the action of selection for quantitative traits, which are often observed to have at least some measurable level of standing additive genetic variance (Hill et al., 2008; Mousseau and Roff, 1987). The second, that demography imposes a drift barrier to the efficacy of selection, has been proposed (Kondrashov, 1995; Lynch, 2025, 2020, 2007; Sung et al., 2012) as a fundamental constraint on adaptation in complex organisms.

Selection is not effective when the fitness effects of an allele fall below the reciprocal of effective population size, or

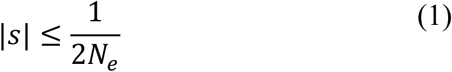

following the scaling of 2 for diploids (Barton, 2022; Kimura, 1983; Ohta, 1973). Below this threshold, termed the drift barrier, the dynamics of an allele are driven primarily by the stochastic force of genetic drift, rather than deterministic fitness effects of selection. This means that deleterious mutations with fitness effects below the drift barrier will fail to be eliminated by selection, and can thus stochastically fix due to drift (Kondrashov, 1995). Concomitantly, beneficial mutations with minute fitness effects will often fail to increase in frequency. Thus, new mutations and already-segregating variants with fitness effects below the drift barrier will evolve in a neutral, or nearly-neutral (Ohta, 2002, 1973), manner.

There is substantial molecular evidence indicating that much of the genome conforms to expectations of near-neutrality (Rockman, 2012). Moreover, mutation rates, which are typically thought be under purifying selection to reduce copy errors, scale negatively with effective population size across taxa (Sung et al., 2012). This suggests that lineages with small effective population size are unable to minimize mutation rate, although alternative explanations based on correlated response to life history diversification can also explain the pattern (Majic et al., 2026). Complex and inefficient cellular physiology in low-*N_e_* metazoans, compared to high-*N_e_* microbes, further suggests wide reaching effects of the drift barrier on adaptation across the tree of life (Lynch, 2020).

These concepts and observations have been invoked to suggest (Lynch, 2025, 2007) that much of the diversity of complex multicellular life may primarily reflect an outcome stochastic non-adaptive processes, rather than adaptation due to natural selection. While developed primarily in the context of explaining molecular traits evolving to a static optimum, the drift barrier hypothesis has been suggested to apply more widely, a point often made in an effort to curb the excesses of adaptationism (Gould and Lewontin, 1979) in the field of evolutionary and molecular biology. Under this drift barrier hypothesis, many features of metazoan life could potentially be viewed as maladaptive baggage that was only able to arise due to the inefficacy of selection at eliminating it.

Even if we are willing to remove our Panglossian blinders, we still can see that this worldview is seemingly at odds with much data from empirical evolutionary ecology. Studies of ongoing adaptation in small populations of metazoans show that natural selection can produce evolutionary response in complex traits in populations with small effective size (Carroll et al., 1997; De Lisle et al., 2022; Grant and Grant, 2006; Losos et al., 1997; Reznick et al., 1997), and that wild populations generally exhibit the capacity to respond to selection (Bonnet et al., 2022). Local adaption (Hereford, 2009), including local adaptation in traits such as physiology (Antonovics, 2006), phenotypic plasticity (Balaguer et al., 2001), and behaviour (Urban, 2007), indicate that small populations can adapt to local optima. At the macroevolutionary scale, parallel or convergent evolution (Bolnick et al., 2018; Losos et al., 1998) – including evolution of phenotypic (De Lisle et al., 2024) and genetic variance (McGlothlin et al., 2022) – in response to replicated environmental gradients indicate that the action of natural selection can produce predictable response that is stable over deep time.

These sorts of studies suggest that much the diversity of metazoan life reflects adaptation in some form, even if evolution is not purely (or even primarily) deterministic. Added to this are a large number of laboratory experimental evolution (Kawecki et al., 2012) and mutation accumulation studies (Katju et al., 2015; McGuigan et al., 2011), demonstrating that adaptation and purging of deleterious alleles in controlled conditions can occur even at low *N_e_*; such evidence does not bear directly on the efficacy of adaptive evolution in the wild, beyond indicating it is likely to be possible.

There are several possible factors that may help to reconcile this apparent conflict between widespread evidence of adaptation in wild metazoans on one hand, and the inevitable constraints on the efficacy of selection in small populations on the other. Natural selection on phenotypes themselves, particularly directional selection, may be strong enough to surpass the drift barrier at the level of at least some segregating alleles or novel mutations. Note that this is difficult to deduce from population genomic data that typically only allows inference of the cumulative action of past selection (Buffalo and Coop, 2020). A recent metanalysis of estimates of *s* revealed that selection can often be strong at the genetic level (Thurman and Barrett, 2016), although nearly all of the approximately 3000 estimates of *s* in this analysis came from two studies (Anderson et al., 2014; Gompert et al., 2014), both of which were manipulative field experiments rather than mensurative studies of selection in unmanipulated populations. Moreover, small effective size, while impeding adaption to a local optimum, will also promote a peak shift to alternative optima (Barton and Charlesworth, 1984; Lande, 1985). Here, I address these potential contributors empirically using a large database of phenotypic selection gradients estimated in wild metazoan populations, as well as a recent compilation of contemporary metazoan effective population sizes. I find that, consistent with theoretical predictions under the drift barrier hypothesis, the strength of directional selection is negatively correlated with effective population size across metazoans. However, analysis of the full distribution of metazoan phenotypic selection and *N_e_* indicates that selection in the wild is often strong enough to overcome the drift barrier across a range of genetic assumptions.

## Results

### Selection-drift balance for quantitative traits

For stabilizing selection acting on a population whose mean is at the optimum, assuming a Gaussian fitness surface given by 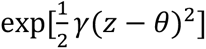, where *γ* is the curvature of the fitness surface (Phillips and Arnold, 1989) and *θ* represents the optimum phenotype, the corresponding threshold mutational effect for selection to be effective is

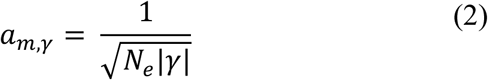

Fixation of a novel deleterious mutation with phenotypic affect *a_m_* results in a displacement of the mean phenotype of the optimum so that *z̄* = *a_m_*, resulting in a phenotypic selection gradient of

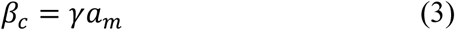

We can thus see that directional phenotypic selection occurring after fixation is stronger than stabilizing selection was on the new mutation, with the corresponding mutational affect at the drift barrier of

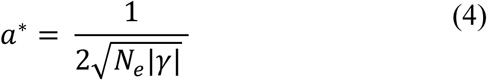

Comparing this to equation 2 illustrates that directional selection resulting from fixation of a deleterious mutation that was at the drift barrier will be effective on mutations twice as small.

We can also see that directional selection should scale with *N_e_* across populations adapting to stationary optima under an Ornstein-Uhlenbeck process (Lande, 1976), where the stationary distribution of the population mean is centred at zero with variance 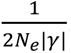. The expected absolute displacement, taken as the mean of the corresponding folded normal distribution, given by 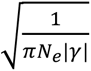. Given equation 3, we can see that the selection gradient is thus

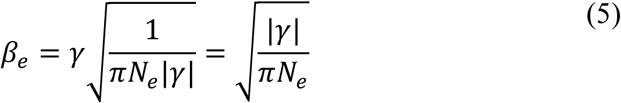

Thus, assuming the curvature of the fitness surface, *γ*, does not vary with *N_e_*, we can see that *β* should scale proportionally to 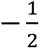 in a log-log regression across lineages, since under equation 5, 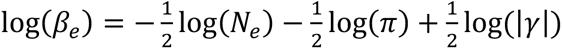.

Using data on the strength of phenotypic selection in wild metazoans (Kingsolver and Diamond, 2011), combined with estimates of metazoan effective population size (Clarke et al., 2024a) we can assess if the absolute strength of directional selection in the wild correlates with *N_e_*. I focus on “contemporary” *N_e_* estimates based on linkage-disequibrium and allele frequency change methods (Clarke et al., 2024a), instead of another recent dataset of eukaryote *N_e_* based on coalescent age (Lewin and Eyre-Walker, 2026), which reflects deep-time effective size and is unrelated to both contemporary measures of evolvability (Abson et al., 2026) and contemporary measures of *N_e_* (Lewin and Eyre-Walker, 2026). Informal metanalysis of 1078 estimates of selection from 21 species and matched median *N_e_* reveals that, indeed, the strength of phenotypic selection is negatively related to log *N_e_* (linear mixed effects model, *t*19 = -2.29, *P* = 0.0335, Figure 1A) and log-log analysis of a reduced dataset (zero estimates of selection dropped) reveals a scaling slope almost quantitatively identical to -1/2 (b = -0.425, SE = 0.23).

**Figure 1.**
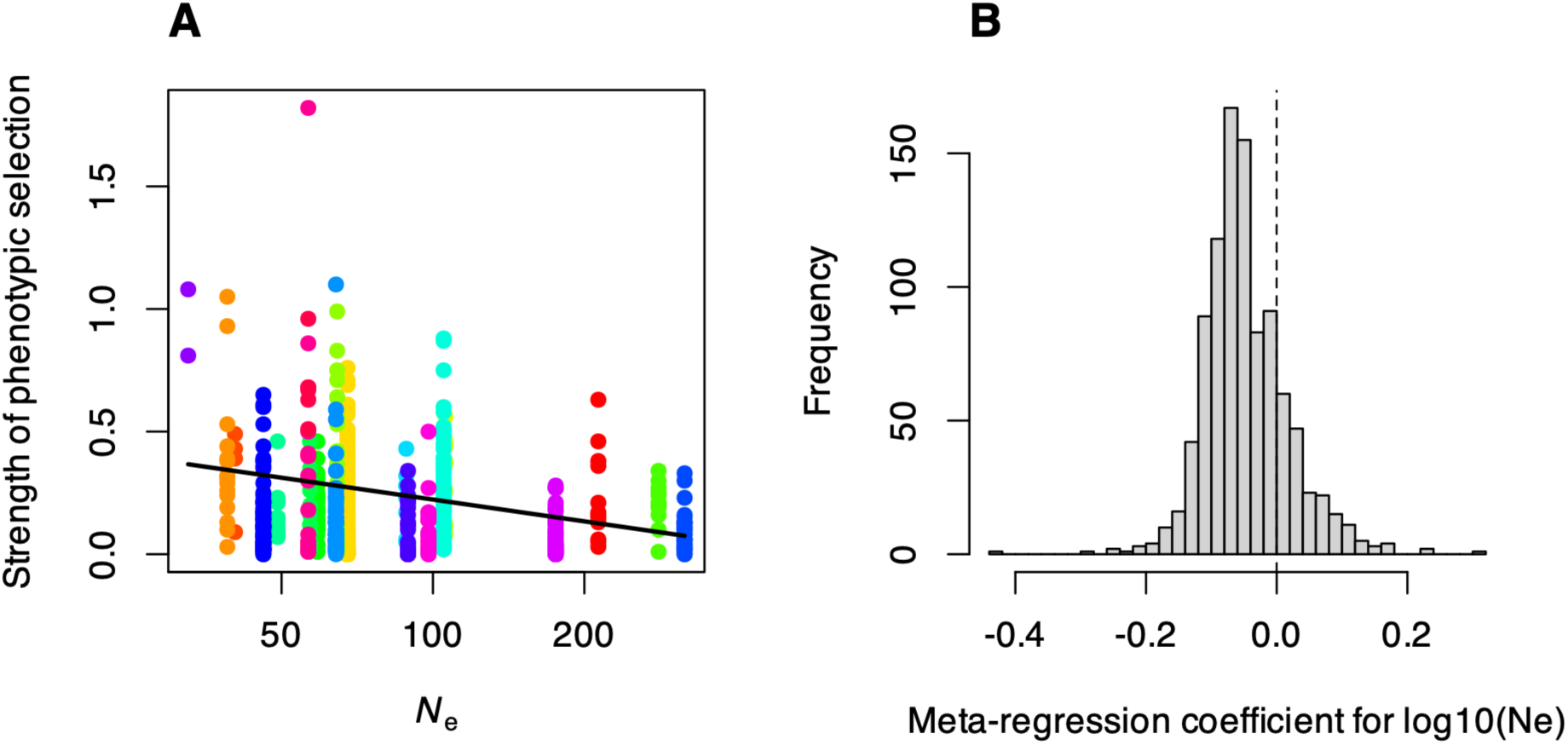
Metaregression shows that directional selection may be stronger in organisms with low effective population size. The absolute strength of phenotypic selection (linear gradients, otherwise differentials if gradients unavailable) is negatively related to the short term estimate of *Ne*. Panel A shows the raw data for all 1078 selection estimates, with the best fit line from a linear mixed model with study as a random effect. The right panel shows the distribution of coefficients from 1000 formal metaregressions on a reduced dataset of 245 estimates for which there was an estimate of the standard error available, where each of the 1000 iterations sampled a different value of *Ne* from the lognormal distribution characterizing the variation in this estimate.

These results hold under formal metanalysis of a reduced dataset of 245 selection estimates from 8 species, for which standard errors for the selection coefficients are available (slope estimate on arithmetic scale: -0.165, SE = 0.085, *P* = 0.0528; estimate on log scale: -0.64, SE = 0.16, *P* < 0.0001). Resampling to propagate variation in estimates of *N_e_* indicates that these results are largely robust to uncertainty in estimates of effective size (Figure 1B).

### The strength of phenotypic selection on standing variance

We can now focus on how the strength of phenotypic selection may translate into selection on underling genetic variants. Importantly, by focusing on phenotypes we can understand contemporary selection on ecologically important traits on the scale of single generations. This contrasts with molecular genetic approaches that typically only allow the inference of the cumulative action of past selection on variants or genomic regions.

We assume *L* biallelic loci with independent additive effects. The relationship between phenotypic selection and the absolute selection coefficient acting on a allele at locus *i* is given by

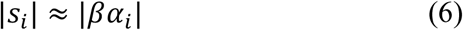

where *α_i_* is the average phenotypic effect of a substitution at locus *i* (Barton and Turelli, 1991). Assuming traits are variance-standardized, the additive genetic variance for the trait contributed by locus *i* can be described as

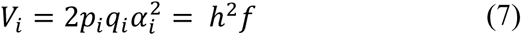

Where *f* is the fraction of the heritability explained by variance at locus *i* and *p* and *q* are the allele frequencies, and summing across loci Σ*V_i_* = *h*^2^. We can see

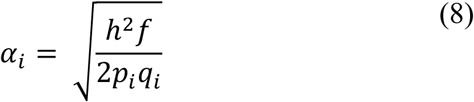

Thus, the strength of selection on underlying loci is a function of the strength of selection on the phenotype (*β*) and how genetic variance is distributed across loci. Under the simple case of even phenotypic effects across all *L* loci and maximum genetic variance at each locus, 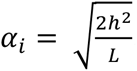. Alternatively, it is possible to sample *f* from a gamma distribution and allele frequencies from a uniform or beta distribution, to explore hypothetical distributions of *s_i_* that would be expected to be generated by selection observed in the wild.

We see from Figure 2A that the strength of selection at underlying loci resulting from ecological phenotypic selection may often be quite strong. For example, even for highly polygenic traits, where *L* = 1000 or 100, we can see that a non-trivial fraction of alleles are expected to fall above the drift barrier even for the case of very small *N_e_*, on the order of 100 (Figure 2B). At *L* = 1000, the mean *si* is 0.004 (standard deviation = 0.011), and at *L* = 100, the mean *si* is 0.013 (standard deviation = 0.036; figure 2A). This is assuming a uniform sampling of minor allele frequencies, and a distribution of phenotypic effects sampled from a gamma distribution with shape parameter equal to 0.2, corresponding to a scenario of many alleles of small effect and a small number of large effect alleles. The general conclusions are robust to alternative assumptions (Supplemental Figures S1-S2), including sampling from a skewed distribution of minor allele frequences (Supplemental Figure S3). Noteworthy is that allowing for more rare alleles, which must have larger phenotypic effects to account for a given fraction of heritability, shifts the expected distribution of *si* to larger values (Supplemental Figure S4). Given that such a skewed minor allele frequency is most consistent with that often observed, the results under more simplistic assumptions (e.g., Figure 2, S1-S3) are conservative.

**Figure 2.**
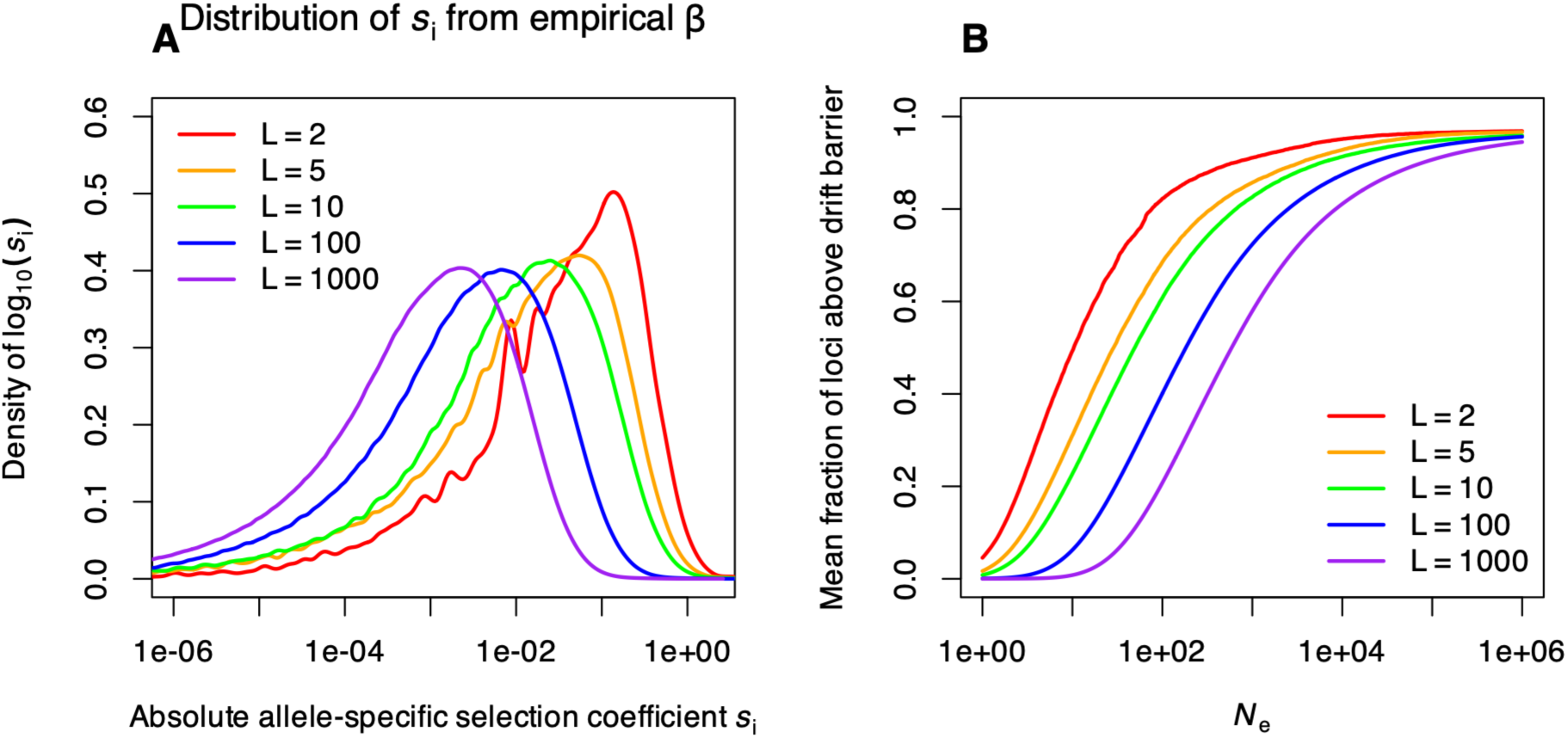
Phenotypic selection in wild populations can generate selection on segregating variants that often exceeds the drift barrier. Panel A shows the distribution of selection coefficients *s*i that is obtained by simulation (see text) from the empirical distribution of phenotypic selection gradients in the Kingsolver and Diamond dataset, for different assumed number of loci *L*. Distribution for each level of *L* is pooled from 100 samples of allelic effects (from a gamma distribution with shape parameter 0.2) and allele frequencies from a uniform distribution. Panel B shows the corresponding fraction of loci for each level of L that is above the drift barrier, as a function of *Ne*.

We can also use the distribution of metazoan contemporary *N_e_* to ask the same question in reverse: Given a value of *N_e_*, what strength of phenotypic selection would be required to result in some significant fraction of underlying alleles to experience effective selection? The distribution of contemporary *N_e_* from Clarke et al. is replotted in Figure 3A. Assuming traits are polygenic (*L* = 100), and the distribution of phenotypic effects of segregating variants is sampled as above, there is a nearly perfect correspondence between the distribution of *β* that would be required to generate selection above the drift barrier for 50% of loci underlying a trait and the observed empirical distribution of *β* from the Kingsolver and Diamond dataset (Figure 3B). Changing *L*, or focusing on a different fraction of loci than the median, has obvious effects on the distribution of *β* required to exceed the drift barrier. Nonetheless, the striking correspondence in Figure 3B supports the analysis above and indicates that, even if realized *N_e_* is remarkably low, phenotypic selection in the wild is often strong enough to be effective.

**Figure 3.**
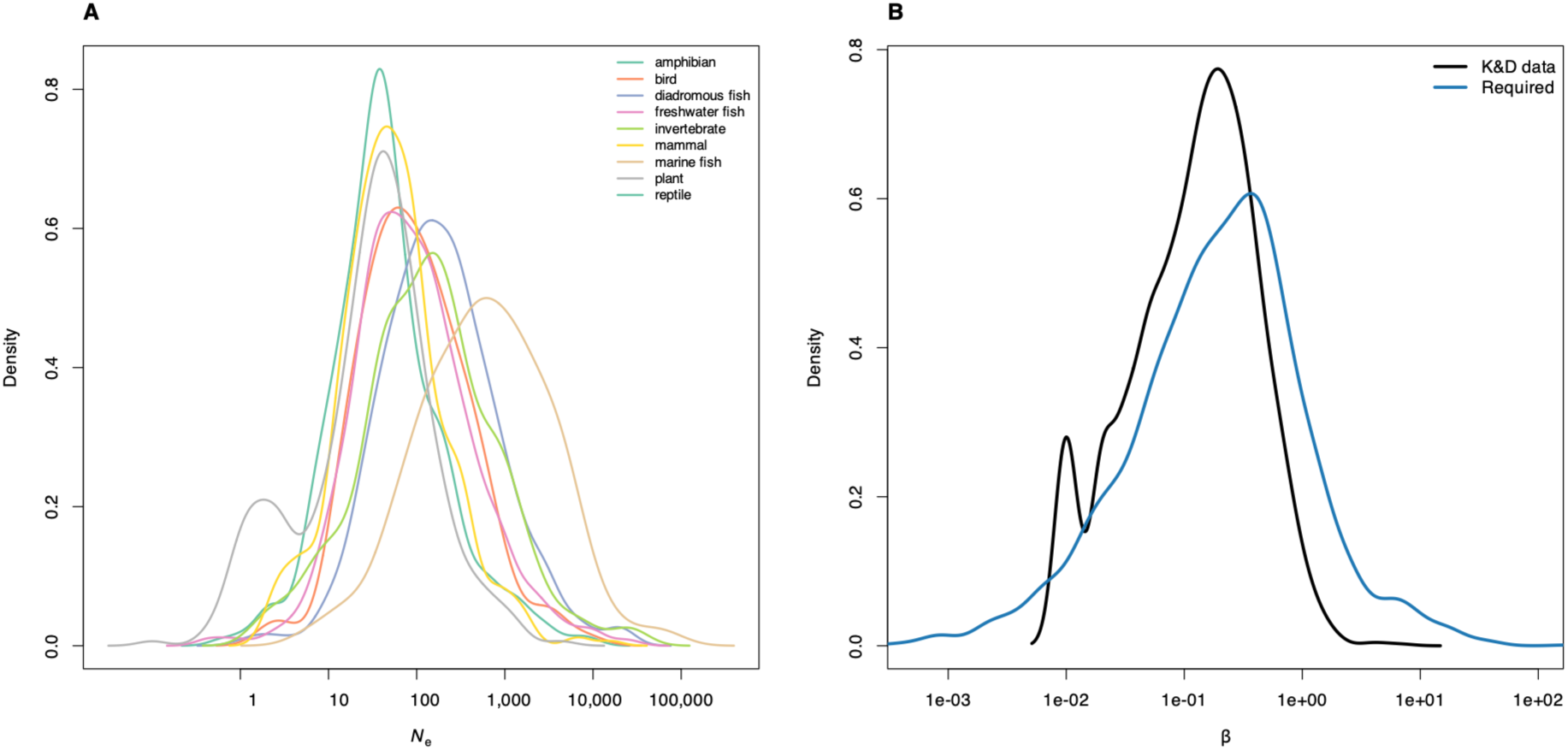
Observed distributions of metazoan Ne and phenotypic selection suggest directional selection is effective at many loci. Panel A shows the kernel density of contemporary metazoan *Ne* estimates from the Clarke et al. 2024 dataset. Panel B shows the corresponding distribution of selection gradients required for the median allelic effect to exceed the drift barrier in blue, averaged over 100 sampled genetic archectures for each observed *Ne*, with allelic effects sampled as in figure 2, and assuming *L* = 100. In black is the kernal density of selection gradients in the Kingsolver and Diamond 2011 dataset.

### The strength of phenotypic selection on novel mutations

Translating a phenotypic selection gradient into the strength of selection expected on a novel mutation is far more straightforward, and less assumption-laden, than translation to selection on already-segregating variants. Given a variance-standardized phenotypic selection gradient *β*, the threshold phenotypic effect (in units of standard deviations) of a mutation that is affected by selection (that is, the magnitude of a mutation that would produce a fitness effect at the drift barrier) is given by

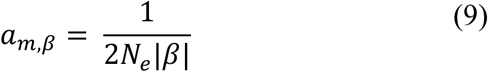

Some insight can be gained into what observable values of *a_m_* may look like from mutation accumulation experiments, which typically measure the per-generation mutational variance *Vm*, which can be scaled to the phenotypic variance. Although *Vm* can vary by orders of magnitude, scaled values of *Vm* on the order of 10^−4^ are typical (Conradsen et al., 2022).

Taking the square root to obtain a per-generation aggregate deviation suggests a per-generation mutational contribution on the order of 0.01 standard deviations are typical in mutation accumulation experiments. As shown in figure 4, for many quantitative traits under directional selection in the wild, selection is often strong enough to exceed the drift barrier for such values. This is consistent with conclusions from analysis of mutational variance that selection is often effective on mutational input (Houle et al., 1996).

**Figure 4.**
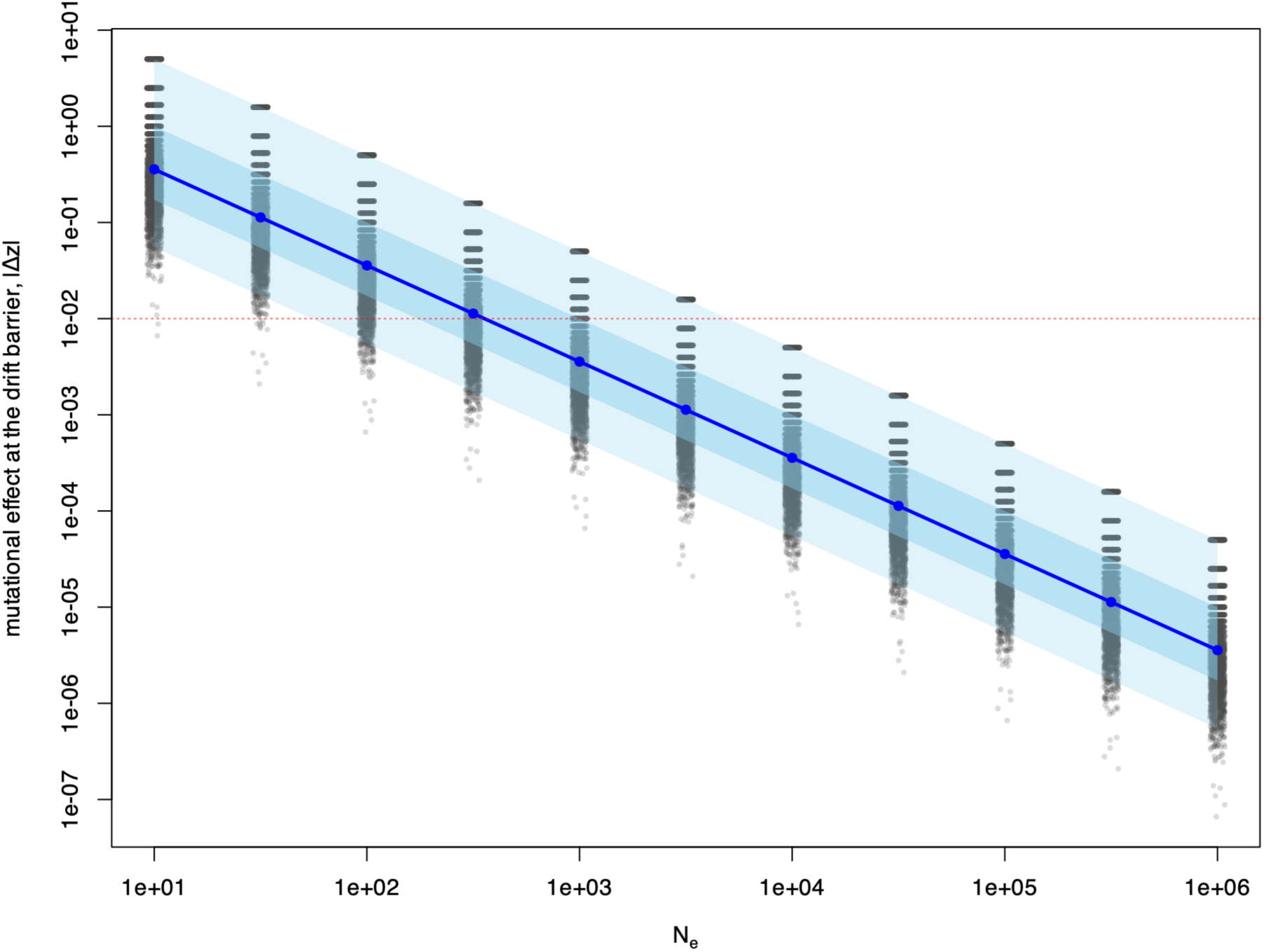
Phenotypic mutational effects required to exceed the drift barrier reveal that selection is often strong enough for selection to be effective on new mutations. The phenotypic effect at the drift barrier is shown for each empirical phenotypic selection gradient in the Kingsolver and Diamond 2011 dataset, shown across a range of *Ne*. Blue line shows the median of the distribution, dark blue the interquartile range, and light blue is the 95% percentile interval.

### Peak shifts induced by genetic drift

From Barton, Charlesworth, and Lande, we can obtain the approximate wait time *t* for a drift-induced shift from the vicinity of on adaptive optimum to another,

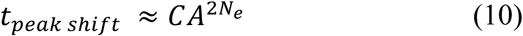

where *C* is a function of genetic variance and the curvature of the topography at the optimum and in the valley, while

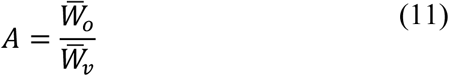

is the ratio of mean fitness at the current optimum relative to mean fitness in the valley (Barton and Charlesworth, 1984; Lande, 1985). Thus, the probability, or rate (Estes and Arnold, 2007), of a drift-induced peak shift can be described as

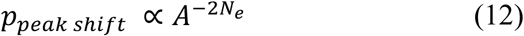

Indicating that peak shifts occur most readily across shallow fitness valleys and at low *Ne*.

Thus, constraints on adaptation imposed by the drift barrier, as well as the likelihood of a peak shift to a new adaptive zone, are both functions of *Ne*. Figure 5A shows this relationship, with the relative probability of a peak shift plotted across a range of *A* values. Noteworthy is that the rate of peak shifts is exponentially related to *Ne*, while the drift barrier is a function of the reciprocal of *Ne*. Thus as *Ne* declines, the drift barrier becomes harder to surmount, yet the likelihood of a peak shift increases more quickly unless the fitness valley is quite deep (Figure 5B).

**Figure 5.**
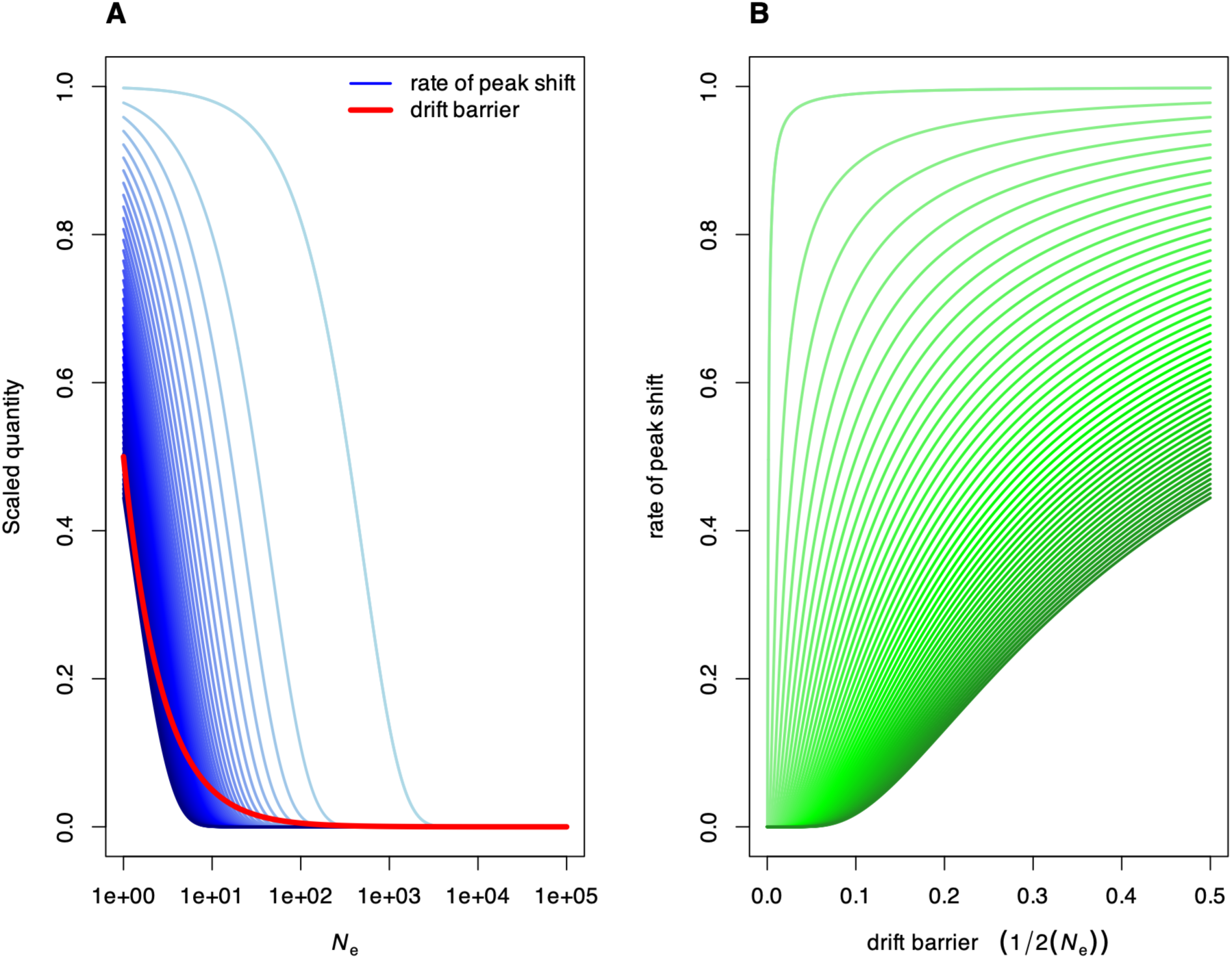
Constraints on adaptive perfection and probability of an adaptive peak shift both scale with effective population size. Panel A shows the drift barrier, 1 / 2*Ne*, and the relative probability of a peak shift by genetic drift, *A^−2Ne^*, across a range of depths of the fitness valley *A* (from 1.0001, light blue, to 1.5, dark blue). Panel B shows the drift barrier and relative probability of a peak shift plotted against each other, with green indicating variation in *A*.

## Discussion

The efficacy of natural selection is fundamentally limited by effective population size, because in small populations the stochastic force of genetic drift will overwhelm weak selection. For many metazoan populations – which may have very small effective sizes – the constraints on adaptation imposed by the drift barrier are thought to be substantial. Yet, for a polygenic trait under stabilizing selection, theory suggest that compensatory directional selection generated by the corresponding fixation of deleterious alleles will be substantially more effective than the initial force generated by stabilizing selection. Here, I find that the strength of phenotypic directional selection in the wild scales with contemporary *Ne* according to a -1/2 relationship, consistent with theoretical expectations under selection-drift balance. I further show that the strength of selection is typically strong enough to overcome the drift barrier, even for quite small *Ne* and highly polygenic assumptions. This same distribution of phenotypic selection gradients indicates selection is strong enough to generate effective selection on novel mutations of realistic magnitude for morphological traits. These results indicate that while drift plays a role in limiting adaptive perfection, the drift barrier may not present a fundamental impediment to the resulting response to directional selection on ecologically important traits. When combined with the diversifying effects of drift-induced peak shifts, which in principle allow finite populations to explore the adaptive landscape, this can potentially reconcile widespread evidence of phenotypic adaptation in metazoans that must also face fundamental constraints on molecular adaptation imposed by the drift barrier.

Quantitative genetic theory predicts a negative scaling relationship between the *absolute* strength of directional selection and *Ne* under selection-drift balance, since small populations will on average deviate more from the optimum under drift even if the expectation for the mean remains at the optimum (Lande, 1976). Strikingly, I found evidence for a negative scaling relationship between the absolute strength of phenotypic selection in wild populations and contemporary *Ne* estimates, with a scaling slope matching the theoretical prediction of -1/2. While this result is based on hundreds of selection estimates, the taxonomic sampling is relatively sparse, and so this should be viewed somewhat preliminarily. Nonetheless, the scaling between phenotypic selection and contemporary *Ne* is consistent with a dual role for selection and drift in governing the dynamics of phenotypic evolution in wild populations. Alternatively, other explanations for such a relationship exist; for example selection itself can reduce *Ne* (Rockwell and Barrowclough, 1995), particularly sexual selection (Nunney, 1993).

While the above result suggests a key role for drift in pushing small populations away from their optima, further analysis of the strength of phenotypic selection suggests that resulting selection is often strong enough to generate some evolutionary response even for small populations and highly polygenic traits. Even for traits whose variance is governed by segregating variants at 1000 or 100 loci with a strongly right-skewed distribution of phenotypic effects, we find a substantial fraction of allele-specific selection coefficients would still be expected to fall above the drift barrier, meaning that we would expect allele frequency change at these loci. However, a substantial fraction of selection coefficients are also observed to fall below the drift barrier in such a case, indicating that the constraints imposed by the drift barrier may nonetheless limit the observed response to selection.

Short term empirical estimates of *Ne* indicate that metazoan populations typically exhibit very low effective size on a generation-to-generation timescale. For example, the median *Ne* across all classes in the Clarke et al. data (Clarke et al., 2024a, 2024b) is 80, with a mean of 534 (Figure 2A). One implication of this observation is that drift-induced peak shifts, especially to local optima and on a more contemporary timescale, may be a more viable contributor to diversification than has been typically appreciated. For example, Estes and Arnold (Estes and Arnold, 2007) rejected the peak shift model as an explanation for deep-time vertebrate body size evolution, primarily on the grounds that their model fits suggested *Ne* between 200-750 was required to explain the data, consistent with Lande’s (Lande, 1985) observation that wait times become exceptional beyond *Ne* on the order of 10^2^. Indeed, deep time estimates of vertebrate *Ne* (Lewin and Eyre-Walker, 2026) suggest that drift-induced peak shifts should essentially never occur. On the other hand, 89% of *Ne* estimates in the Clarke et al. data fall below 750, 71% below 200, and 91% are on or below the order of 10^2^. Thus, conclusions on the role of drift-induced peak shifts depends fundamentally on whether the relevant (for the dynamics of selection and phenotypic evolution) scale of *Ne* is recent genetic change or coalescent age. My analysis of contemporary phenotypic selection suggests a role for “contemporary” *Ne*, as does another recent study linking contemporary selection and *Ne* to long term evolution in *Drosophila* (Brändén and De Lisle, 2026), and short-term *Ne* has also been proposed to play a role in explaining Lewontin’s paradox of genetic variation (Buffalo, 2021). Others have argued that deep time *Ne* estimates are more likely to be relevant for understanding evolvability and longer term evolutionary dynamics, although these estimates do not correlate strongly with evolvability (Abson et al., 2026). There is clearly some need to reconcile the disparity in various *Ne* estimates and when each may apply (Wang, 2005).

Focusing on contemporary *Ne* estimates makes conclusions on the limitations imposed by the drift barrier conservative, as these estimates are orders of magnitude lower than coalescent *Ne*. For example, the lowest estimate for a vertebrate *Ne* in a recent dataset of coalescent-scale *Ne* (Lewin and Eyre-Walker, 2026) was 12500, for the green monkey *Chlorocebus sabaeus*, with metazoan *Ne* estimates ranging from 10^4^-10^6^. As can be seen from Figures 2 and 4, such large effective sizes pose little barrier to directional selection of the magnitude often observed in the wild, regardless of the genetic underpinnings of phenotypic variation.

Using estimates of directional phenotypic selection in the wild to obtain the distribution of *s* has the distinct advantage in that it provides insight into the potential strength of selection that could occur on genetic variants in actual unmanipulated wild populations on the timescale of a single generation. Of course, the estimates of phenotypic selection are themselves imperfect, and suffer from a number of well-documented limitations such as the problem of missing traits (Morrissey et al., 2010), component fitness measures, and missing fractions of the phenotypic distribution (Mittell et al., 2025). Nonetheless, phenotypic selection is far easier to directly measure than selection on regions of the genome, especially on ecological timescales, and so we have a far more complete understanding it.

Most quantitative traits are thought to be highly polygenic (Barton, 2022; Sella and Barton, 2019), with important consequence for molecular and phenotypic evolution. Polygenic traits may evolve and respond to selection even if the signatures of such selection at the molecular level are minute, and thus challenging or impossible to detect (Rockman, 2012). Additive polygenic variation means that fixation of deleterious alleles can potentially be compensated for by selection on alleles at other loci (Barton, 2022). Consistent with this, my analysis suggests that selection on the level of phenotypes in the wild is often strong enough surpass the drift barrier at some loci, while many others would be expected to evolve in a nearly-neutral manner. One implication is that, for polygenic traits, we expect segregating variants of large effect to more often surpass the drift barrier and thus consistently respond to selection in a manner detectable by population genomic methods giving the impression that trait evolution is largely governed by few alleles of major-effect (Rockman, 2012). Similarly, in declining or small populations, we expect selection to have greatest efficacy on variants with unconditional fitness effects (Connallon et al., 2025).

Despite a shared foundation (Fisher, 1930), there remain significant theoretical and empirical divides between population and quantitative genetics (Walsh and Lynch, 2018). While quantitative genetic analysis of phenotypic selection (Lande and Arnold, 1983; Svensson, 2023) focuses on temporary covariance between traits and fitness, molecular population genetic approaches rely on signatures of persistent longer-term molecular change to infer selection (Hahn, 2019), although some short term estimates of *s* do exist (Thurman and Barrett, 2016). Rapid response to strong selection on polygenic traits can often result in weak or undetectable molecular signatures of a selective sweep (Barghi et al., 2020; Thornton, 2019). Even if phenotypic selection is strong enough to produce allele frequency change at loci of small phenotypic effect, it may be the case that these changes are likely to be largely undetectable (Rockman, 2012).

The drift barrier hypothesis has been used to argue that much of the diversity of complex life may be an outcome of non-adaptive processes (Lynch, 2025, 2007), with the primary evidence coming from molecular genetic data (Lynch, 2020; Sung et al., 2012). I have shown that predictions of selection-drift balance also hold when confronted with quantitative genetic estimates of contemporary selection and demography, but that directional selection may nonetheless often be strong enough to partially overcome the drift barrier. Together these results indicate a dual role for adaptation and chance in governing the dynamics of contemporary phenotypic evolution in metazoans.

## Materials and Methods

### Data acquisition

I obtained 4693 variance-standardized estimates of directional selection from the database of Kingsolver and Diamond (https://doi.org/10.5061/dryad.7996) (Kingsolver and Diamond, 2011). These are estimates of phenotypic selection in wild populations of 114 species of plants and animals, and reflect the typical strength of selection observed in ecological studies of natural selection in nature. Estimates of metazoan effective population sizes have been recently compiled by Clarke et al. (Clarke et al., 2024a), (data at https://doi.org/10.5061/DRYAD.P2NGF1VZM), with positive finite estimates from 3576 populations from 596 species. Both of these datasets contain multiple estimates of each parameter (selection or *Ne*) from multiple populations/studies of each species, and for selection, multiple traits.

### Meta regression of Ne and selection

Selection and effective size were matched by species or genus across the two datasets, with the assistance of GPT 5.6. Ten species with selection estimates (either gradients, or if they were unavailable, differentials) had corresponding *Ne* estimates available, while 11 others had estimates from at least one congener. All positive finite matched *Ne* estimates were retained (e.g., all congener estimates for a species in the selection dataset), and from these the median, mean, and standard deviation of *Ne* was computed. Thus, the final matched dataset contained 1080 estimates of phenotypic selection from 21 species with corresponding measures of *Ne* and its variation. Two exceptional values of selection, above 4, were excluded as outliers, although their inclusion did not change results. Excluding measures of selection that did not have standard errors yielded 245 estimates of selection from 8 species. Given this massive reduction, I proceeded with both informal metanalysis with the full dataset of 1080 (1078 excluding outliers) dataset, as well as formal metanalysis with the reduced dataset of 245 estimates.

For the informal metanalysis, I fitted a mixed model with the absolute strength of selection as the response and the median *Ne* as a fixed effect. Median *Ne* was log10 transformed. The model included a random effect for selection study nested within species. I repeated this analysis with log10 transformed absolute selection, noting that some estimates recorded as zero had to be dropped. For formal meta analysis, I used the metafor package in R to fit a meta regression model with absolute strength of selection as the response, the square of the standard error as a measure of variance in the observations, log10 median *Ne* as a moderator (fixed effect), and selection study and genus as random effects. I repeated this in log-log space after dropping estimated selection with an exact recorded value of zero, using the delta method to transform the standard errors. In order to understand the effects of propagating variation in *Ne*, I then repeated the meta regression sampling (1000x) *Ne* from a lognormal distribution centred on the original median *Ne* with standard deviation of the lognormal calculated from the original mean and standard deviation. For this I used all of the data possible, i.e. treating selection on the arithmetic scale. For each sampling run, I refitted the same meta regression model described above.

### Analysis of the distribution of selection gradients

For calculation of selection on underlying genetic variants starting from phenotypic selection estimates, I focus on analysis of selection gradients, *β*, for which there are 2819 estimates in the Kingsolver and Diamond database. Given an estimate of |*β*|, the resulting strength of selection on allele at a biallelic locus *i* requires assumptions about the number of loci, as well as 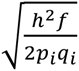, where *h*^2^ is the heritability of the trait, the fraction *f* of the variance contributed by locus *i* (noting that ∑*_L_ f*=1), and the allele frequencies at the locus. I assumed *h*^2^ = 0.3, which is conservative and in line with typical estimates of heritability of quantitative traits, and explored a range of *L* from 2 to 1000. *f* and *p_i_q_i_* where treated as stochastic. I sampled *f* from a gamma distribution with shape parameter of .2, which corresponds to a strongly skewed distribution where a few loci contribute substantially to heritability, but most contribute small effects. Samples were scaled to sum to unity (that is, generating Dirichlet random variables for *f*). I also explored sampling from a uniform distribution, as well as a various values of shape parameter for the gamma distribution. For allele frequencies, I sampled a uniform distribution with minor allele frequency bounded at a minimum of 0.05 or 0.001, as well as a beta distribution with shape parameters of 0.3, 3. I also tried the alternative of assuming the maximum variance at each locus, a fixed *p_i_* = 0.5. These various alternatives had little effect on conclusions, with the main contribution being the assumption of number of loci *L*. For each value of *L* in the figures, I sampled 100 genetic architectures.

All analyses were performed in R (version 4.5.3); GPT 5.6 was used as a coding aid.

## Author Contributions

S. De Lisle performed all aspects of this research

## Classification

Biological sciences, evolutionary biology

## Acknowledgements

I thank E. Svensson for comments on the manuscript, and L. Rowe, J. Stinchcombe, and the evolutionary ecology group at Karlstad University for discussion and encouragement. Funding was provided by the Swedish Research Council (Vetanskapsrådet grants no. 2024-04599), Formas (grant no. 2021-01096), and KK-stiftelsen (grant no. 2024STG).

## Supplemental Figures

**Figure S1.**
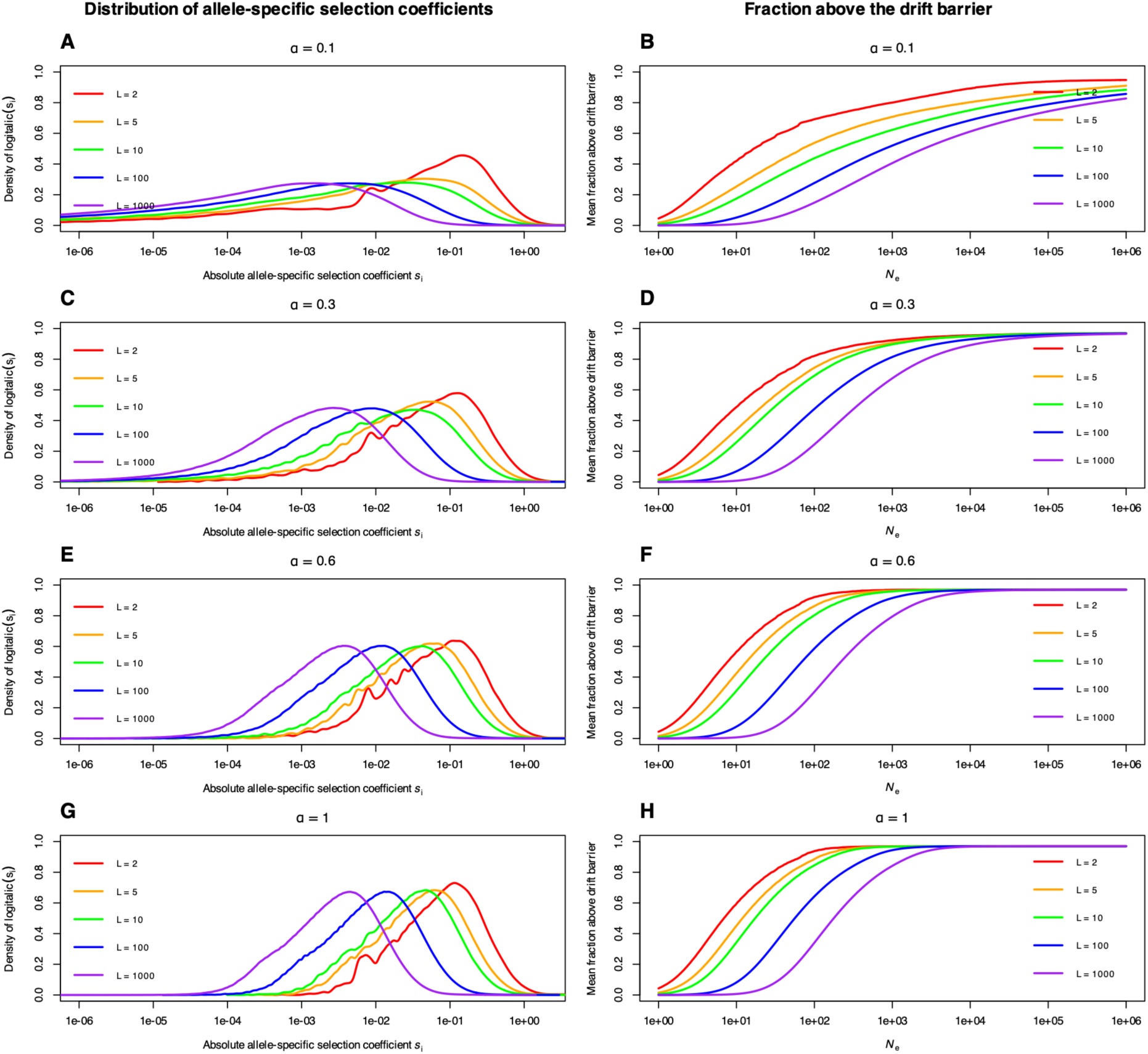
Allele-specific selection coefficients calculated from phenotypic gradients under varying assumptions about the distribution of allelic effects. Each row repeats Figure 2 under a different value of the gamma shape parameter α from which heritability fractions are sampled. Other details, including allele frequency sampling, are as described for Figure 2.

**Figure S2.**
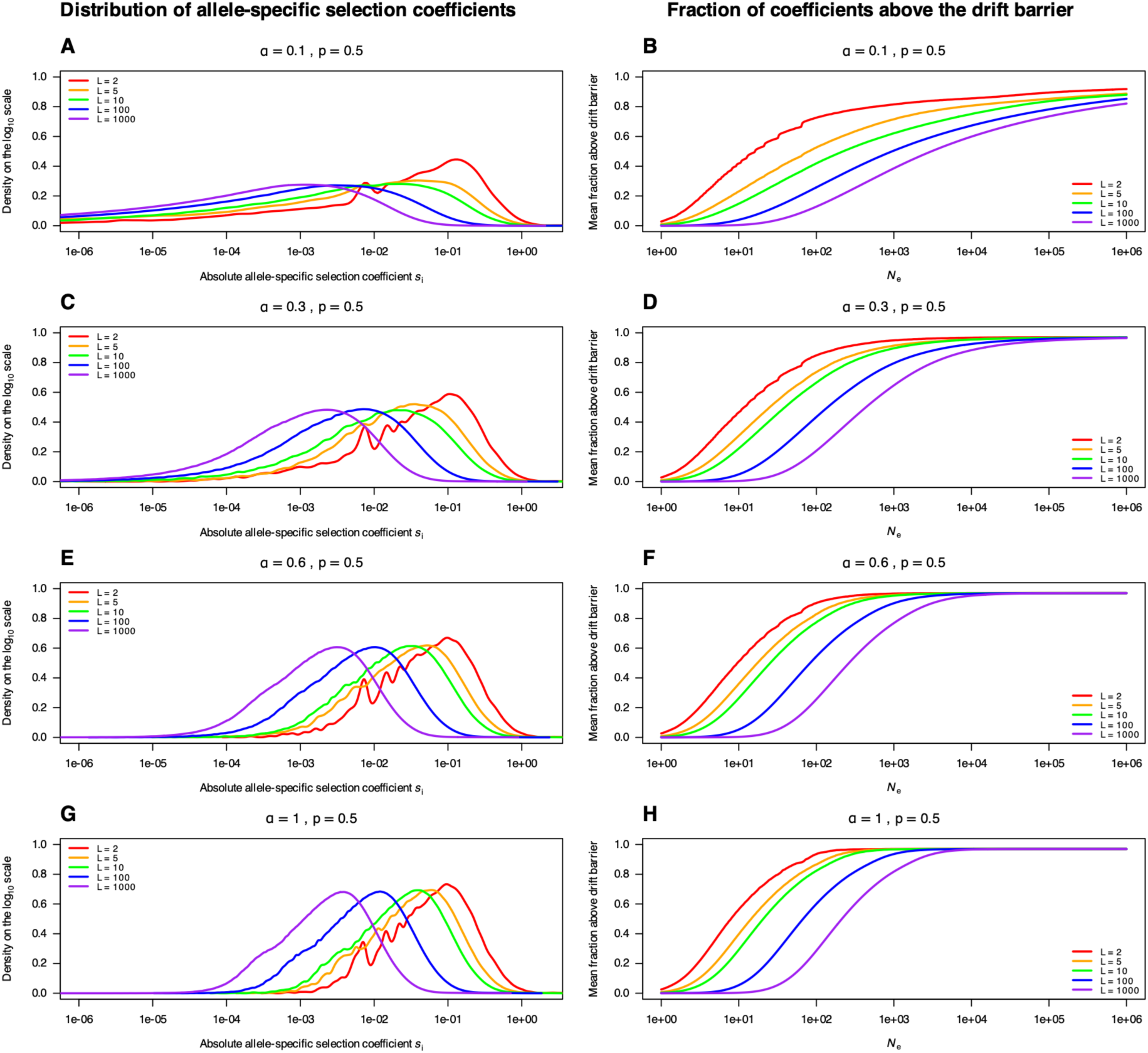
Allele-specific selection coefficients calculated from phenotypic gradients under varying assumptions about the distribution of allelic effects assuming fixed allele frequences. Each row repeats Figure 2 under a different value of the gamma shape parameter α from which heritability fractions are sampled, and assuming a fixed allele frequency of 0.5 across all loci. Other details are as described for Figure 2.

**Figure S3.**
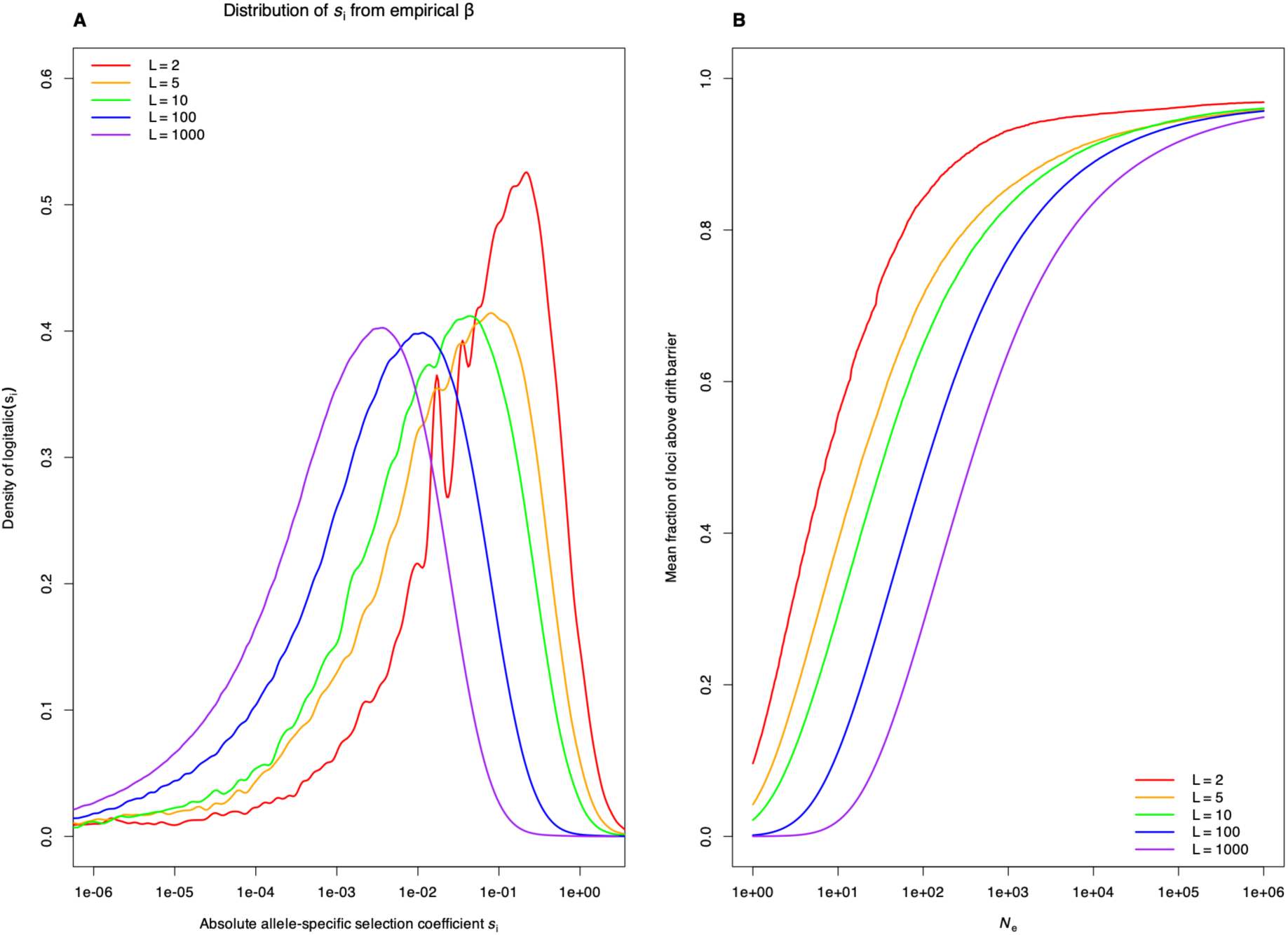
Allele-specific selection coefficients calculated from phenotypic gradients, when sampling minor allele frequences from a beta distribution. Assuming α = 0.3 and β = 3, with a lower cutoff of MAF = 0.05. Other details, including allele frequency sampling, are as described for Figure 2.

**Figure S4.**
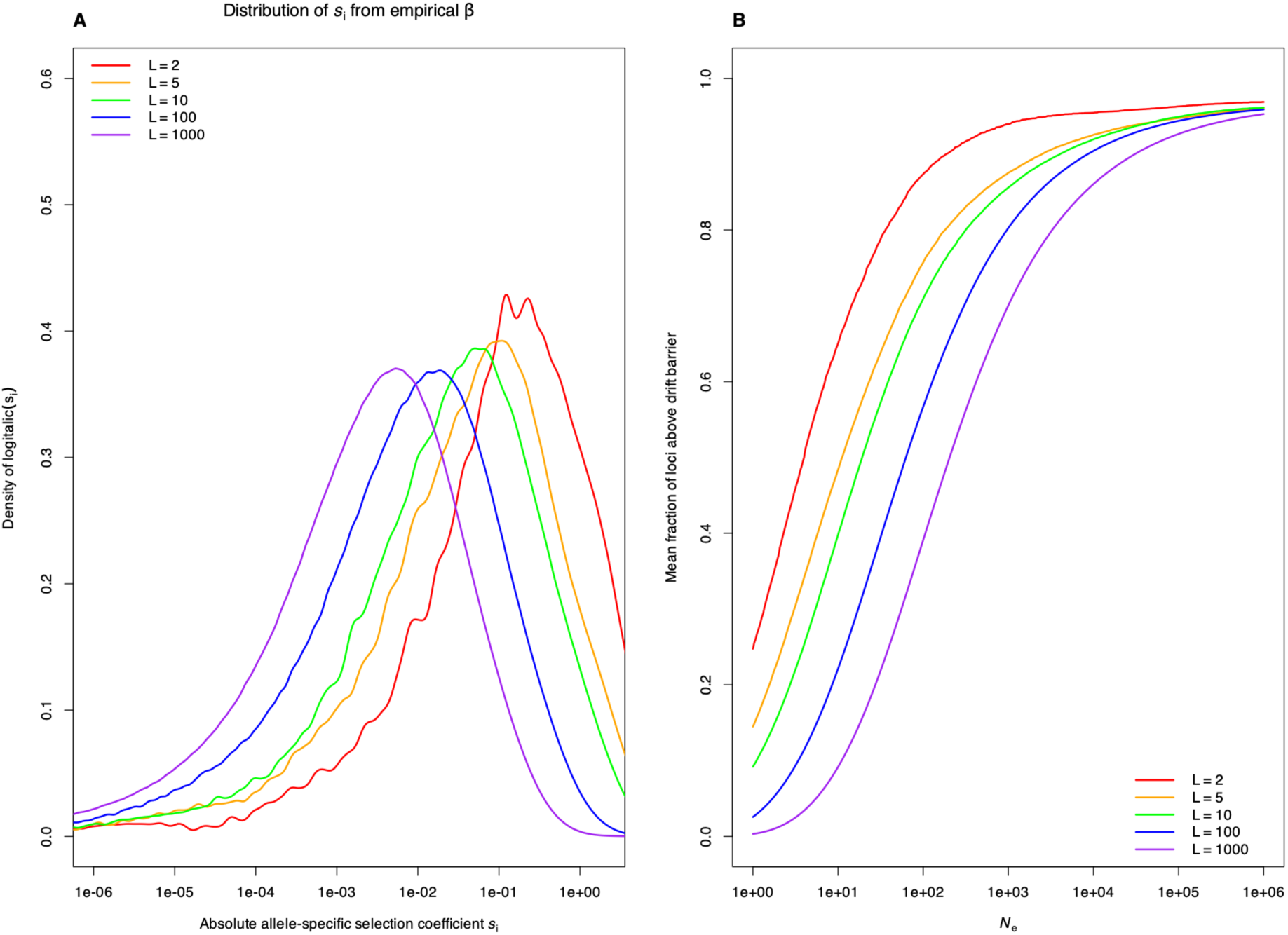
Allele-specific selection coefficients calculated from phenotypic gradients, when sampling minor allele frequences from a beta distribution. Assuming α = 0.3 and β = 3, with a lower cutoff of MAF = 0.001. Other details, including allele frequency sampling, are as described for Figure 2.

